# Impact of light pollution at night on male reproductive success in Japanese medaka (*Oryzias latipes*)

**DOI:** 10.1101/2023.04.17.536935

**Authors:** Lauren E. Closs, Muhammad Rahmad Royan, Amin Sayyari, Ian Mayer, Finn-Arne Weltzien, Dianne M. Baker, Romain Fontaine

## Abstract

Environmental light is perceived and anticipated by organisms to synchronize their biological cycles. Therefore, exposure to artificial light at night could disrupt diurnal and seasonal rhythmicity. Reproduction is a complex physiological process involving integration of environmental signals by the brain, and release of endocrine signals by the pituitary that regulate gametogenesis and spawning. In addition, males from many species form a dominance hierarchy that, through a combination of aggressive and protective behavior, influences their reproductive success. In this study, we investigated the effect of different light regimes, including light pollution at night and continuous light, on the fitness of male fish within a dominance hierarchy using a model fish, the Japanese medaka. In normal light/dark rhythm conditions, we observed that dominant males are more aggressive, remain closer to the female, and spend ten-fold more time spawning than subordinates. By using males with different genotypes, we determined the paternity of the progeny and found that even though subordinate males spend less time with the females, they are equally successful at fertilizing eggs in normal light conditions due to an efficient sneaking behavior. However, when exposed to light at night, dominant males fertilize more eggs. We indeed found that when exposed to nocturnal light pollution, dominant males produce higher quality sperm than subordinate males. Surprisingly, we did not find differences in circulating sex steroid levels, pituitary gonadotropin levels, or gonadosomatic index between dominant and subordinate males, neither in control nor night light condition. Continuous light was found to completely inhibit establishment of male hierarchy. This study is the first to report an effect of light pollution on sperm quality with an impact on male fertilization success in any vertebrate. It has broad implications for fish ecology in urban areas with potential impacts on the genetic diversity of these fish populations.

**GRAPHICAL ABSTRACT:** 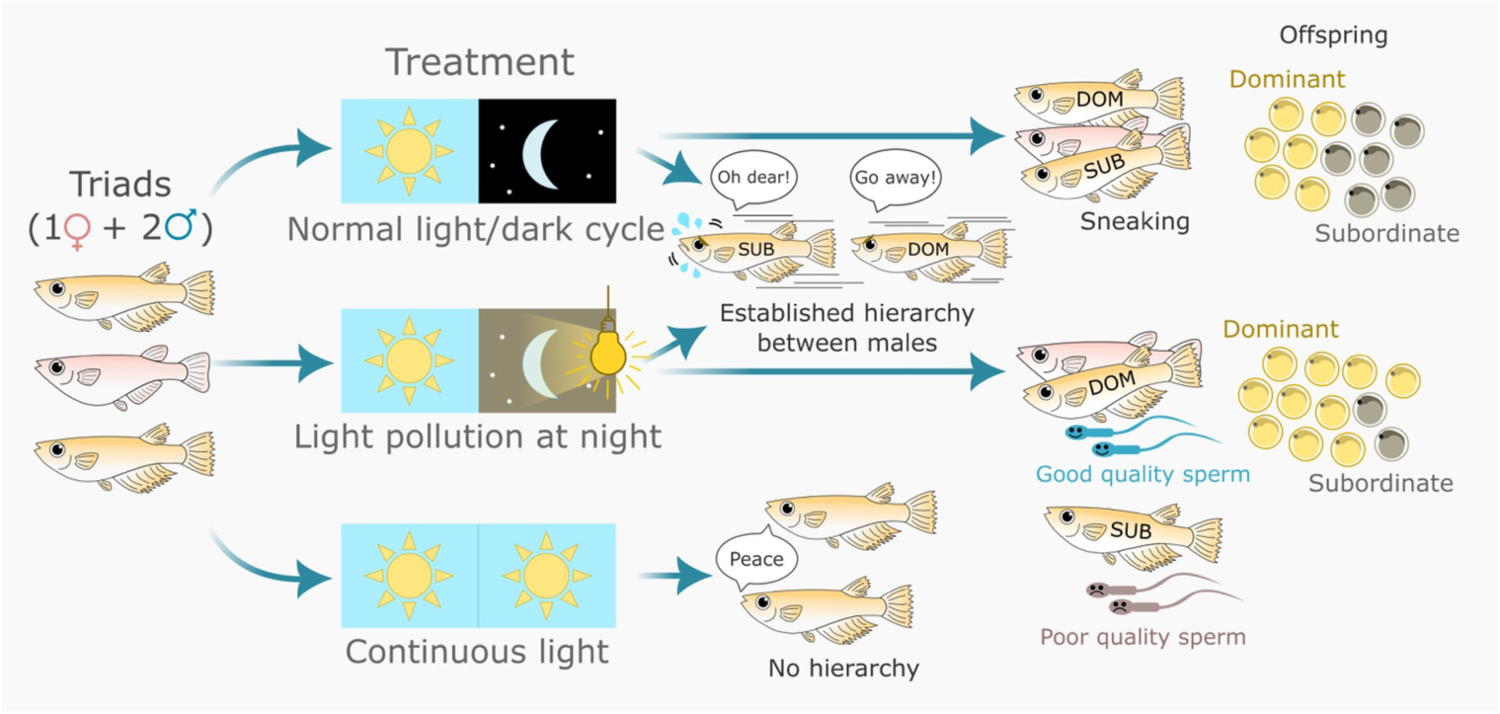

**Highlights:**

- In a triadic relationship, subordinate medaka show sneaking behavior during female spawning, allowing them to produce about 50% of the offspring.
- Continuous light inhibits establishment of male hierarchy.
- Light pollution at night shifts the balance of reproductive efficiency between dominant and subordinate males, benefiting dominant males.
- Exposure to light pollution at night improves sperm quality in dominant fish relative to subordinates, but does not clearly affect reproductive hormone levels.

## INTRODUCTION

Animal behavior and social interactions can influence reproductive fitness. Indeed, position in a dominance hierarchy is sometimes associated with higher reproductive fitness. This is for instance the case in some primate groups where high rank in the hierarchy allows greater reproductive success by increasing breeding opportunities (Alberts et al., 2006). Moreover, through social behaviors, dominant individuals can suppress the activity of the brain-pituitary-gonad (BPG) axis in subordinates (Altmann et al., 1995; Bennett, 1994; Koyama & Kamimura, 2000). These social phenotypes are plastic and quickly reversible, meaning that individual males may switch between dominant and subordinate status multiple times within a lifetime (Fernald & Hirata, 1977). However, although observations in African cichlids revealed a link between dominant status, Lh and steroid levels (Maruska & Fernald, 2013), very little is known about the cellular and molecular mechanisms involved in this dominance hierarchy phenomenon.

As in other vertebrates, sexual maturation and reproduction in fish are controlled by the brain-pituitary-gonad (BPG) axis (Weltzien et al., 2004). In this axis, the two pituitary gonadotropins, luteinizing hormone (Lh) and follicle-stimulating hormone (Fsh), play a central role by regulating gametogenesis and steroidogenesis in the gonads. Lh and Fsh hormone production and secretion are regulated by the brain which integrates environmental signals (Dufour et al., 2010; Yaron et al.; Zohar et al., 2010) and by feedback from the gonads via sex steroids (Romain Fontaine, Muhammad Rahmad Royan, et al., 2020). This axis, mainly through the gonadal sex steroids, also plays a role in the control of reproductive behaviors in all vertebrates, including both mammals (Rhees et al., 1997) and fish (Munakata & Kobayashi, 2009). As such, the BPG axis is the key endocrine axis in the control of vertebrate reproduction, including in the timing of gametogenesis and spawning.

It is widely recognized that photoperiod (daily duration of the light period) is the most important environmental cue in the control of seasonal rhythms in all vertebrates, including in teleost fishes (Rowan, 1938). Accumulating evidence now shows that light pollution from artificial light at night (ALAN), which is present in urbanized areas, can interfere with the photoperiod signal. By increasing the duration of daily light exposure, ALAN dysregulates major physiological functions in many species (Walker II et al., 2019). Currently, about 23% of the Earth is exposed to bright skies at night and the distribution of ALAN is expanding (Kyba et al., 2017). ALAN not only perturbs humans (Lucassen et al., 2016; Navara & Nelson, 2007) and other terrestrial animals (Schroer & Hölker, 2017), but it also affects aquatic habitats (Marangoni et al., 2022). Indeed, at least 22% of coastal regions are exposed to ALAN (Davies et al., 2014), and because more than 50% of the world’s population live within 3 km from a freshwater body (Kummu et al., 2011), a large portion of the freshwater habitats (river, lakes, etc.) are exposed to ALAN.

Surprisingly, in teleost fish, which represent the largest and most diverse group of vertebrates, very little is known about the effects of light pollution, but some recent studies have revealed physiological and behavior impacts. For instance, light pollution at night was found to modify natural circadian cycles of activity and metabolic rate in a coastal fish, the Baunco (*Girella laevifrons*) (Pulgar et al., 2019). Surgeonfish (*Acanthurus triostegus*) larvae exposed to ALAN also displayed changes in behavior, with higher susceptibility to nocturnal predation (O’Connor et al., 2019). In addition, although surgeonfish exposed to ALAN grew faster and heavier, they also experienced significantly higher mortality rates than control fish. Light pollution has also been found to affect fish reproduction and fitness. In a study of smallmouth bass (*Micropterus dolomieu*), ALAN altered the behavior and activity during the nest guarding period (Foster et al., 2016), thus potentially impacting the ability to protect the eggs. ALAN also decreased pituitary gonadotropin and sex steroids levels in European perch (*Perca fluviatilis*) and roach (*Rutilus rutilus*) (Brüning, Kloas, et al., 2018), which could disrupt seasonal reproduction. However, it is unknown whether light pollution affects reproductive fitness by perturbation of reproductive behavior and social interactions.

In addition to light pollution affecting wild fish, light is commonly manipulated in aquaculture to promote growth or to control specific physiological processes. This is for instance the case for farmed Atlantic salmon for which continuous light (Light:Dark (LD) 24:0) is well known to increase growth and to reduce maturation time (Endal et al., 2000; Fjelldal et al., 2018; Oppedal et al., 2006; Stefansson et al., 1991). Interestingly, while it is widely used in aquaculture, whether continuous light (which can be considered as an extreme form of ALAN) affects reproductive behaviors and social interactions also remains unknown.

The Japanese medaka (*Oryzias latipes*) is a powerful model organism to study vertebrate development, physiology, and genetics (for review: (Shima & Mitani; Wittbrodt et al., 2002)). Medaka are small teleost fish widely used in research due to their small size and short generation time which makes their maintenance in a laboratory relatively inexpensive. Due to their wide use globally, numerous tools have been developed and are now available to study this species, such as a sequenced genome (Kasahara et al., 2007) and several transgenic lines with specific cell types, including gonadotropes (Hildahl et al., 2012; Hodne et al., 2019), labeled by fluorescent reporter proteins. Contrary to the other popular small teleost model, the zebrafish (*Danio rerio*), medaka has a genetic sex determination system (Matsuda, 2005; Nanda et al., 2002) like mammals, which has also made them useful for sex determination and sexual dimorphism studies. As an oviparous species, medaka produce transparent eggs that are fertilized externally, and remain attached to the female for few hours. Medaka are daily breeders in laboratory conditions (Fukamachi et al., 2009; Kobayashi et al., 2012; Ono & Uematsu, 1957). Importantly, medaka reproduce seasonally, and reproduction is tightly regulated by photoperiod (Koger et al., 1999; M. R. Royan et al., 2023).

Interestingly, when placed in a triadic relationship (two males with one female), in most cases Japanese medaka quickly form a hierarchy in which the dominant male exhibits a robust mate-guarding behavior, consisting of courtship displays directed toward the female and aggression toward the subordinate male (Yokoi et al., 2015). The effect of dominant status on reproductive success is unclear. While Yokoi and colleagues (Yokoi et al., 2015) observed that dominant males fertilized over 93% of the offspring, by reducing the rival males’ access to potential mating partners, another study reported sneaking behavior by subordinate males during spawning, allowing the ‘sneaker’ males to fertilize 5 - 45% of the eggs in a given clutch (Weir, 2013).

Therefore, in the present study we used the Japanese medaka as a model to investigate the role of pituitary and gonadal factors in the dominance hierarchy and reproductive success in males. We also investigated the effect of nocturnal light pollution, as well as continuous light, on male reproductive fitness and pituitary-gonadal factors.

## MATERIALS AND METHODS

### Animals and light regimes

Adult (5- to 8-month old) Japanese medaka (HdrR strain) were reared in a re-circulating water system (28 °C; pH 7,6) on a LD 14:10 cycle with light turning on at 8 am. Fish were fed three times daily with a combination of dry feed (Zebrafeed, Sparos) and live *Artemia salina*. Sex determination was based on secondary sexual characteristics (Kinoshita et al., 2009). Experiments were conducted in accordance with recommendations on experimental animal welfare at the Norwegian University of Life Sciences.

### Hierarchy behavioral setup

Hierarchical status of males within a triadic relationship (two males co-housed with one female) in 10-L plastic tanks was determined by visual screening. As medaka spawn in the morning soon after the light turns on, fish behavior was recorded using a Nikon D800 camera from when the light turned on until the female spawned. The 10 minutes of video just before the female spawned were analyzed to measure three parameters: 1- protective (mate guarding) behavior, 2- aggressive behavior toward the other male, 3- amount of time spent spawning. For the protective behavior (supplemental video 1 (Closs et al., 2023)), we recorded the tank from above for 10 minutes with only 1-2 cm of water to prevent vertical swimming (figure 1A). We took a picture every minute from the video and measured the distance of each male to the female using ImageJ. The dominant male will actively display mate guarding, (as previously described by (Yokoi et al., 2015) by positioning itself between the female and the subordinate male. For the other two parameters, the video was recorded from the side of the tank. We manually counted the number of fights and “chase with hit” (respectively supplemental videos 2 and 3 (Closs et al., 2023)) as aggressive behavior during the 10 minutes prior to the female spawning. We measured the time spent spawning by each male, as defined by body contractions while alongside females (supplemental video 4 (Closs et al., 2023)), while the female was releasing eggs, as well as any premature spawning attempts during those 10 minutes.

**Figure 1:**
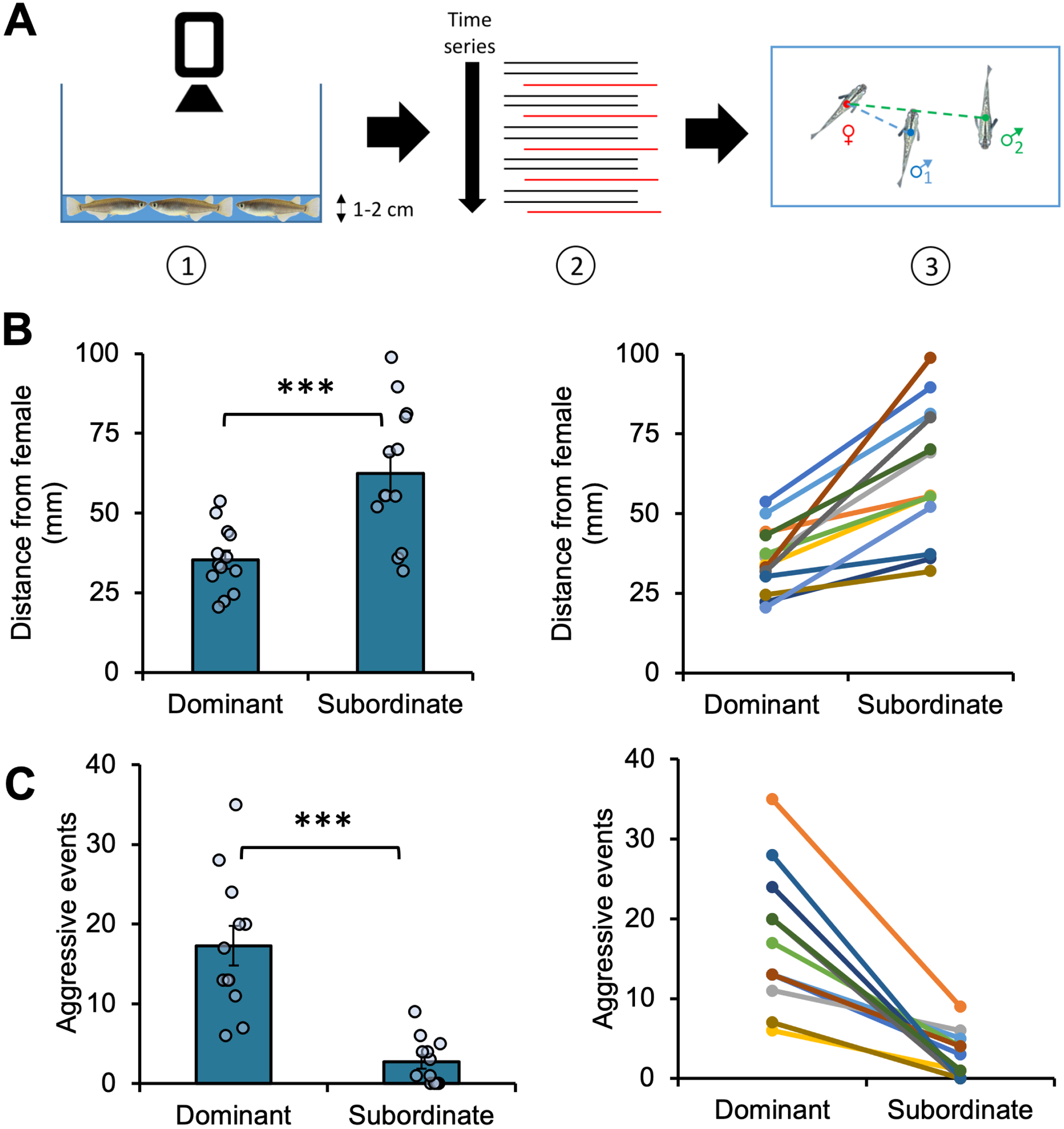
**(A)** Schematic drawing on how the distance between the two males and the female was measured. (1) 10 minutes video were recorded from the top of the tank. The tank was filled only with 1-2 cm of water to limit vertical swimming. (2) Over the series of pictures form the video represented by the horizontal bars, one picture (red) was extracted every minute from the video for (3) measuring the distance between the fish. Dominance parameters in medaka include **(B)** mate guarding behavior, measured as distance of each male from the female (n=13) and **(C)** aggression, counted as aggressive events (n=12). The data are represented as mean ± SEM (left) and graphed according to paired competing males (right). Differences between means were assessed using paired Student’s (B) and Welch’s (C) t-tests, and significant differences are indicated by asterisks (*:p< 0,05; **:p< 0,01; ***:p< 0,001).

### Light condition treatment

Fish were housed on a rack system in a light cabinet with either control light (LD 14:10 with light onset at 8 am), light pollution (LD 14:10+ALAN), or continuous light (LD 24:0) and allowed to acclimatize for two weeks. This time was found sufficient to allow medaka to physiologically adapt to a new photoperiod regime by starting or stopping reproduction following exposure to respectively long or short photoperiod (M. R. Royan et al., 2023). Light levels (Table 1) were measured in two different ways with a digital lux meter (TES-1337, TES Electrical Electronic Corp) and a quantum light pollution sensor (SQ-640, Apogee) at different places within the fish tank. Fish were then set into triads in the light cabinet for 2 more weeks (one to determine the male dominance hierarchy followed by a second to determine the reproductive success).

**Table 1:**
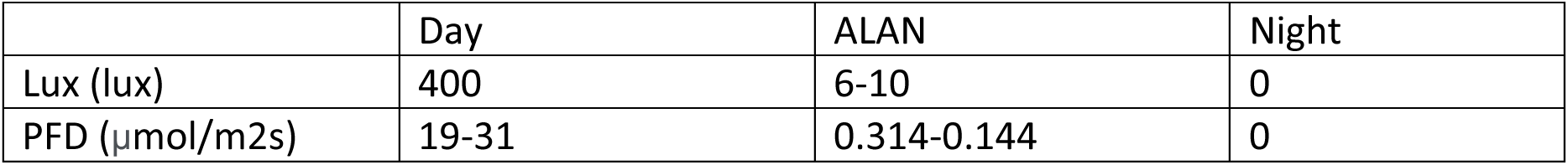
Light intensities used for our experimental conditions.

|  | Day | ALAN | Night |
| --- | --- | --- | --- |
| Lux (lux) | 400 | 6-10 | 0 |
| PFD ($\mu\text{mol}/\text{m}^2\text{s}$) | 19-31 | 0.314-0.144 | 0 |

### Behavior Assay

For each triad, two males of the same age were size-matched and co-housed in a 3-L plastic tank with one female. The males were distinguished by randomly clipping either the top or bottom corner of the tail fin (supplemental figure 1A) under anesthesia with 0.03% Tricaine methanesulfonate (MS222, Sigma). The behavior of the fish in triads was observed for 1-3 minutes each at three time points in the morning for 5-8 days. The male exhibiting the dominant behavior, characterized as guarding the female and aggression toward the other male, was identified for each triad and recorded. All observations were blind, without knowledge of the records from previous time points. Males that exhibited aggressive behavior and close proximity with the female for more than 80% of the time analyzed were considered dominant, while the other males in the triads were considered subordinate.

To measure the activity (time spent in different zones in the water column and distance swum) of the fish exposed to different light regimes, water column tests (n= 2 for each treatment group) were performed by video recording for 10 minutes with a camera (acA1300-60gm Basler). This test involved recording the time individual fish (2 males and 1 female) spent in each of three different zones (shown in figure 5B), as measured automatically using Ethovision software (Noldus).

To determine if fish activity (time spent moving and velocity) differed between treatment groups during nighttime and daytime, the activity of individual fish placed in triads was recorded over a 24-hour period using infrared backlight panels (Noldus) (for night recording) and a Basler camera (n= 4 for each treatment groups). Then, data from 8:00-9:00 and midnight-6:00 were analyzed using the Ethovision software with default parameters to measure average velocity and average time spent moving during the morning and night intervals.

### Measuring Reproductive Success

Reproductive success of dominant and subordinate males under control light conditions (14:10) and ALAN (14-:10+ALAN) was measured using triads consisting of one homozygous transgenic male tg(*lhb:hrGfP-II*) (Hildahl et al., 2012), one wild-type male, and one wild-type female. For the control condition, n= 17 triads, and for ALAN, n= 15 triads. After the behavior assay and determination of dominance and subordinance, 20-30 embryos were collected from 2-4 clutches for each female. The presence or lack of GFP signal in the gut (supplemental figure 2) was visually screened at 48-h post-fertilization under a fluorescent microscope to determine paternity by the transgenic or wild type male. All screenings were done blind and only fertile fish were used.

This experiment was also replicated twice under control light conditions, once using the same technique of GFP screening (n= 10), and a second time using triads consisting of one homozygous tg(*lhb:hrGfp-II)* male, one homozygous tg(*fshb:dsRed2*) male (Hodne et al., 2019), and one wild-type female (n= 10). Following the behavior assay and determination of dominance and subordinance hierarchy, we collected at least 20 embryos and incubated them for 1-5 days in Petri dishes at 26 °C. Embryo DNA was extracted directly from crushed embryos using the Phire Animal Tissue Direct PCR kit (Thermo Scientific). To determine paternity, individual embryos were genotyped by PCR for both hrGfp-II and ds-Red2, followed by gel electrophoresis in 2% agarose gels. Sequences for the PCR primers are shown in Table 2. Reproductive success was determined by comparing the percentage of eggs fertilized by dominant males versus subordinate males.

**Table 2.**
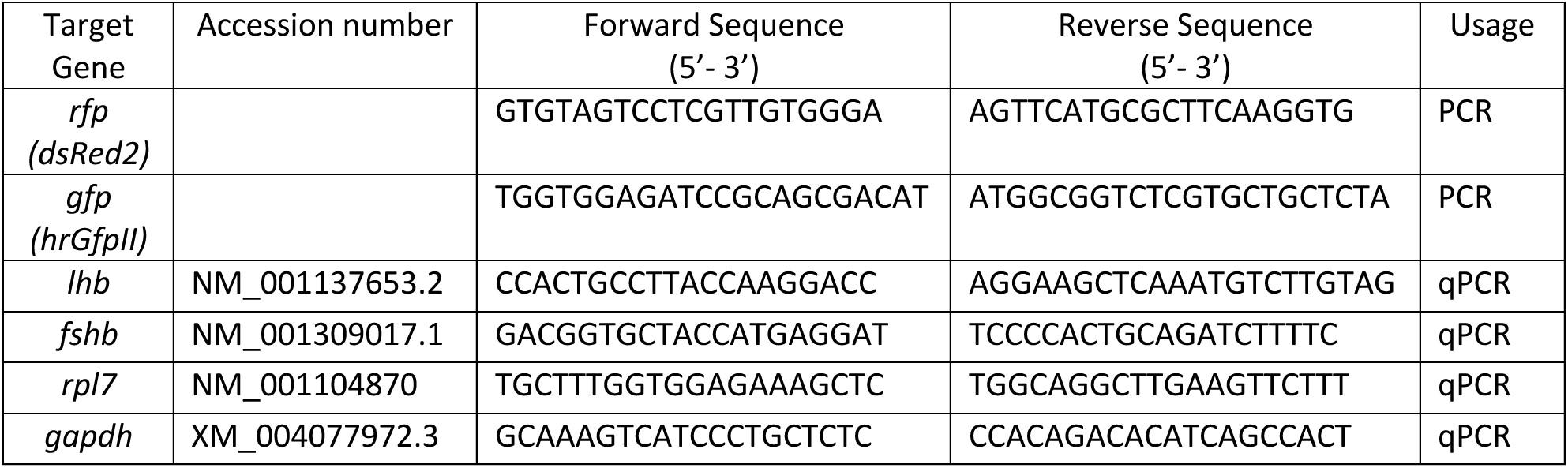
Primer sequences for PCR genotyping and qPCR.

### Sample collection

Dominant and subordinate males were euthanized with an overdose of MS222 and weighed. Blood was collected from the peduncle vein using a glass capillary as described previously (Royan et al., 2020). The blood was stored at −80 °C for later measurement of steroid concentrations. Then, the testes were excised as previously described (Closs et al., 2022) and quickly weighed to calculate gonadosomatic index (GSI= 100 x (gonad weight/fish weight)) before the sperm was extracted by homogenizing the testes in 120 μL Hank’s balanced salt solution (HBSS, 287 mOsmol/kg, Sigma) with forceps. Extracts were stored on ice until sperm quality was assessed. Finally, pituitaries were isolated as described previously (Ager-Wick et al., 2018), transferred to 300 μL of Trizol (Thermo Scientific), and stored at −80 °C for later quantification of gonadotropin mRNA levels. All samples were collected during the nighttime, between 23:00 and 1:00 when sex steroids and gonadotropin mRNA levels are high in medaka (Koger et al., 1999; Muhammad Rahmad Royan et al., 2023).

### Assessment of sperm quality parameters

Sperm quality parameters, including percent motile sperm, progressivity, and velocity, were analyzed using a computer-assisted sperm analysis (CASA) approach. We used the SCA® (Sperm Class Analyzer® CASA System-MICROPTIC) 1-h after collection as previously described in details (Closs et al., 2022), using disposable counting chamber slides (2-chamber counting slides; 20 µm nominal chamber depth, Leja Products B.V., Nieuw-Vennep) and an Eclipse Ts2R microscope (Nikon).

### Quantitative polymerase chain reaction (qPCR)

qPCR and data analysis were performed as previously described (Burow et al., 2019; Royan et al., 2021). Total pituitary mRNA was extracted by homogenizing pituitaries in Trizol with 6 lysing beads (MP Biomedicals) with a tabletop vortex. RNA concentration and purity were measured using an Epoch microplate spectrophotometer (Biotek). Total RNA (214 ng) was reverse transcribed using SuperScript III reverse transcriptase (Invitrogen) and 5 µM random hexamer primers (Invitrogen) following the manufacturer’s instructions. qPCR was performed using LightCycler 480 SYBR Green I Master (Roche). Each cDNA sample was measured in duplicate using 3 μL of 4x diluted cDNA per 10 μL reaction in a LightCycler96 Instrument (Roche). qPCR parameters were set to 10 minutes of preincubation at 95 °C, 42 cycles of 95 °C for 10 seconds, 60 °C for 10 seconds, and 72 °C for 15 seconds, followed by melting curve analysis to evaluate PCR product specificity. Cq values were normalized using the combination of housekeeping genes *rpl7* and *gapdh*. Primer sequences for each gene used are displayed in Table 2.

### Sex steroid extraction and enzyme-linked immunosorbent assay (ELISA)

Steroids were extracted from 1 μL of blood using diethyl ether (Muhammad Rahmad Royan et al., 2023). Once dry, the pellet was resuspended in 50 μL of ELISA buffer. Testosterone (T) and 11-Ketotestosterone (11-KT) ELISA kits (Cayman Chemicals) were used to measure T and 11-KT. Cortisol concentrations were measured using a Cortisol ELISA kit (Arbor Assays). Sex steroid concentrations were calculated with a standard curve fitted by a 4-parameter logistic regression (R^2^ > 0.99).

### Statistics

All statistics were performed using either Jamovi or GraphPad Prism. For the time spent spawning by males before and during female oviposition, the Rout test was used with aggressive parameters (Q=0,1%) to remove definitive outliers. Normality tests were performed using Shapiro-Wilk tests prior to running parametric tests; non-parametric tests were used in cases of non-normal distributions as stated in figure legends. Significance was set to P<0.05.

## RESULTS

### Dominant males are more aggressive and spend more time spawning but do not have higher reproductive fitness

When two males were grouped with one female, a hierarchy was clearly established with one male, which we later designated as the dominant male, exhibiting strong guarding behavior as shown by the significantly reduced average distance from the female, and more aggressivity (figure 1B,C). We then confirmed that clipping of the bottom or top corner of the tail fin (supplemental figure 1A) did not affect the hierarchy outcome (supplemental figure 1B) and that dominant and subordinate males did not differ in mean body mass or size (supplemental figure 1C,D).

Dominant males spent 10-fold more time displaying spawning behavior than the subordinate males (figure 2A). We further investigated the spawning time for each male by distinguishing the time spent with females before and during oviposition. We found that dominant males spent a considerable time contracting their body alongside females prior to egg release (figure 2B), suggesting that they may release sperm before the eggs are released. This was not the case for subordinate males.

**Figure 2:**
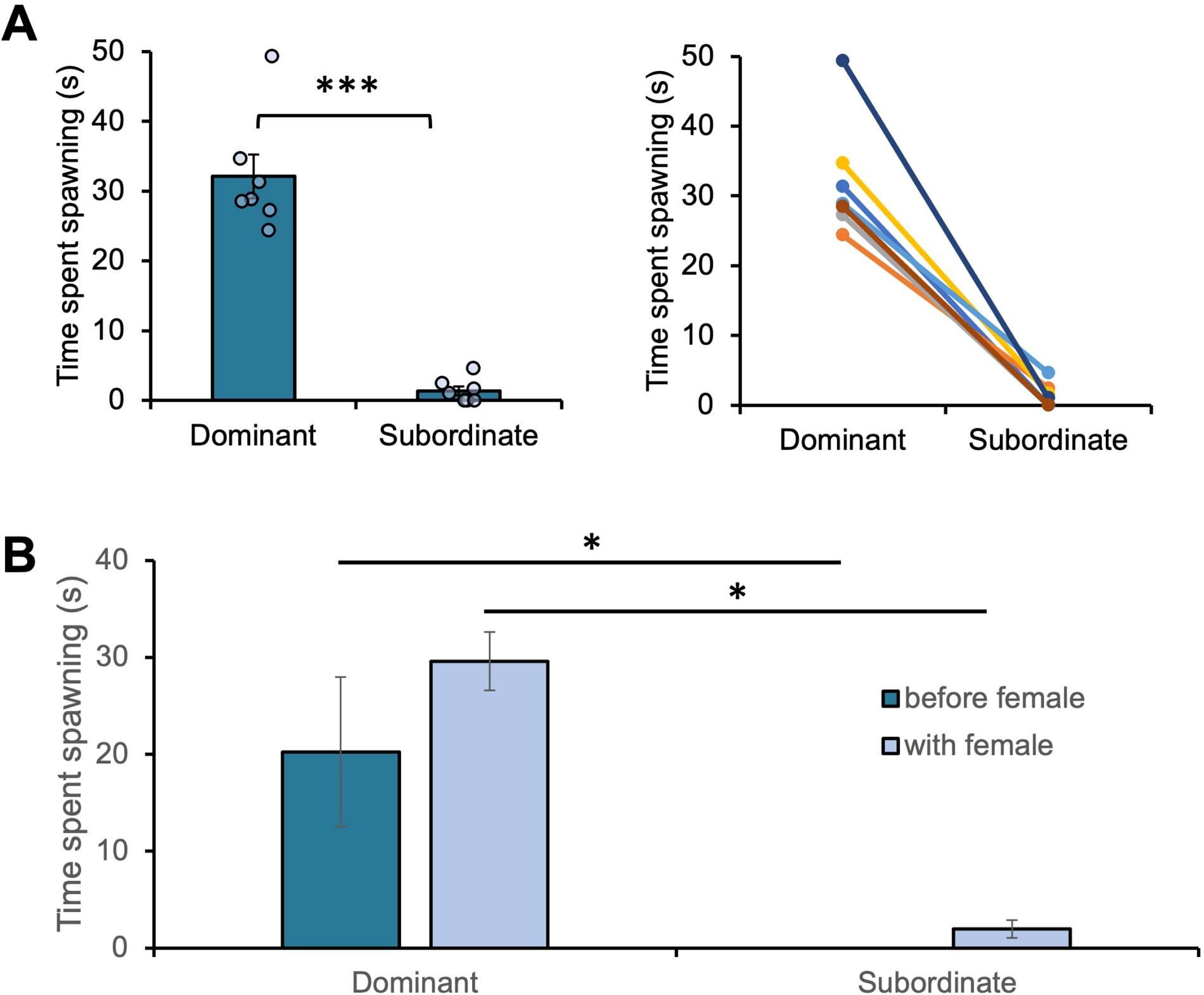
Another parameter that could be used to identifiy dominant males is **(A)** the time spent spawning (n=7). The data are represented as mean ± SEM (left) and graphed according to paired competing males (right). The time spawning was then subdivided **(B)** in before and during female oviposition (n= 7-9). The data are represented as mean ± SEM. Differences between means were assessed using paired Student’s t test (A) and Kruskal-Wallis test (B), and significant differences are indicated by asterisks (*:p< 0,05; **:p< 0,01; ***:p< 0,001).

Interestingly, when looking at the paternity of the embryos using either visual screening of Gfp (figure 3A) or genotyping (data not shown), we found that subordinates are as likely as dominant males to fertilize eggs and produce progeny. Indeed, subordinate males fertilize on average 48.4% ± 7.88 (mean ±SEM) of eggs while dominants fertilize 51.6% ± 7.88. We confirmed that neither the genotype of the fish nor clipping the bottom or top corner of the tail fin affected the progeny origin (supplemental figure 1E,F). Together, these results suggest that dominant males, by starting to spawn before the eggs are released by the females, may have lower sperm quantity than subordinate males at the time the female is releasing eggs. This would allow subordinates males to fertilize about half of the eggs despite the limited access to females.

**Figure 3:**
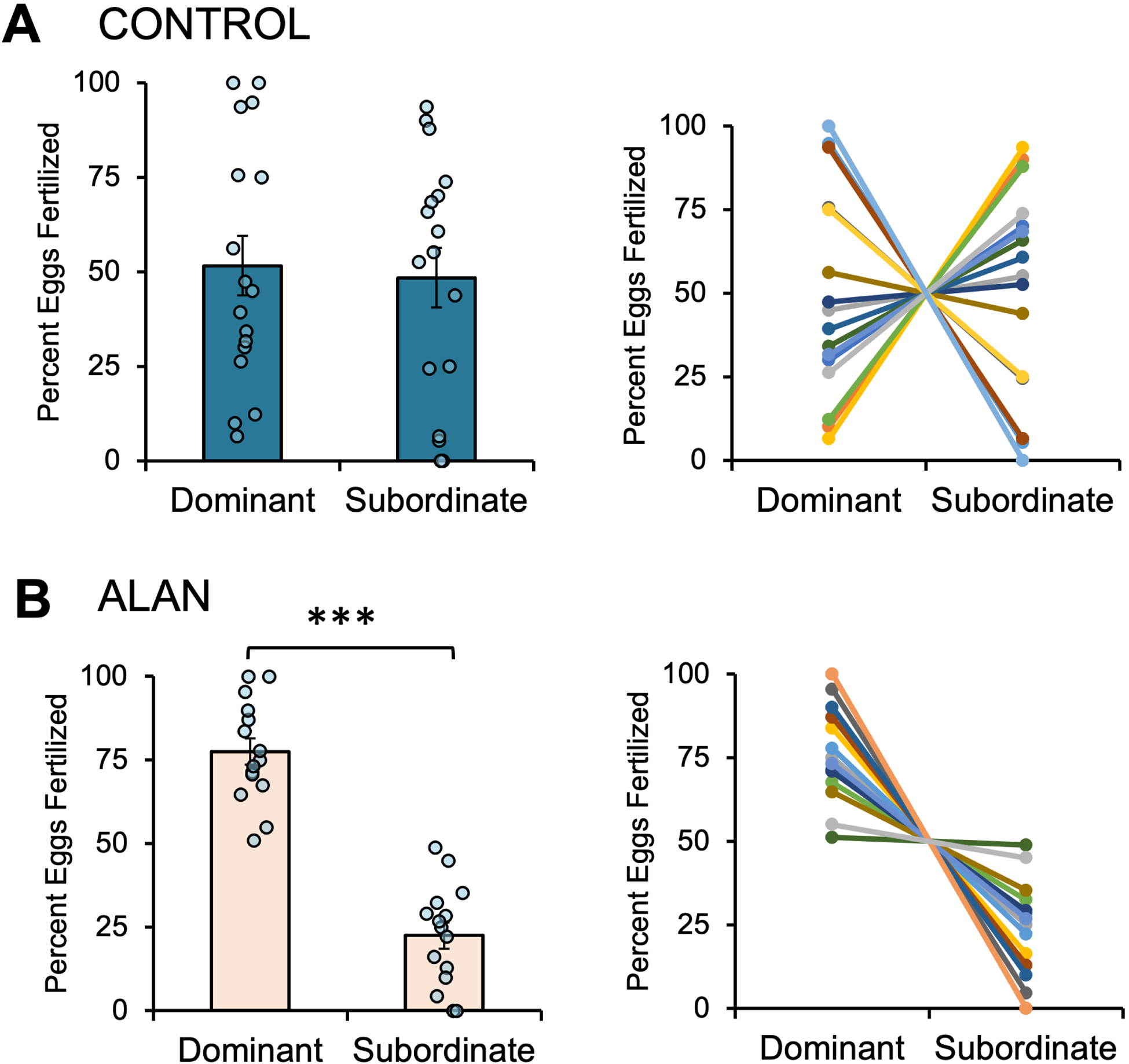
Paternity of the progeniture determined by the percentage of eggs fertilized by the dominant and subordinate males exposed to **(A)** control light (n=17) and **(B)** artificial light at night (n=15). The data are represented as mean ± SEM (left) and graphed according to paired competing males (right). Differences between means were assessed using paired Student’s t-tests, and significant differences are indicated by asterisks (*:p< 0,05; **:p< 0,01; ***:p< 0,001).

### Light pollution influences the reproductive success of males

We then tested the effect of different light regimes on male hierarchy, behavior, and reproductive success. While we found that ALAN had no visible effect on male hierarchy, with dominant males still easily identifiable, ALAN surprisingly resulted in changes to the paternity ratio of offspring (figure 3B). Dominant fish exposed to light pollution fertilized significantly more eggs (77.5% ± 3.9), while subordinates fertilized only 22.5% ± 3.9. The dominant males tended to guard females more closely than the subordinate males (figure 4A), but mean distance did not significantly differ. In addition, the dominant males showed significantly higher aggressivity and spent more time spawning compared to the subordinate males (figure 4B,C). We thus investigated whether ALAN affected the time males spent spawning before and during female oviposition. We found that dominant males exposed to ALAN spent significantly more time spawning before and during oviposition than subordinate males (figure 4D), a pattern similar to that seen with control light (figure 2B). Although some subordinate males showed spawning behavior before female oviposition in ALAN condition (figure 4D), which was not the case under control light conditions (figure 2B), the change was not significantly different (P = 0,48 with a Mann-Whitney test), suggesting that the shift in fitness is not due to altered reproductive behavior.

**Figure 4:**
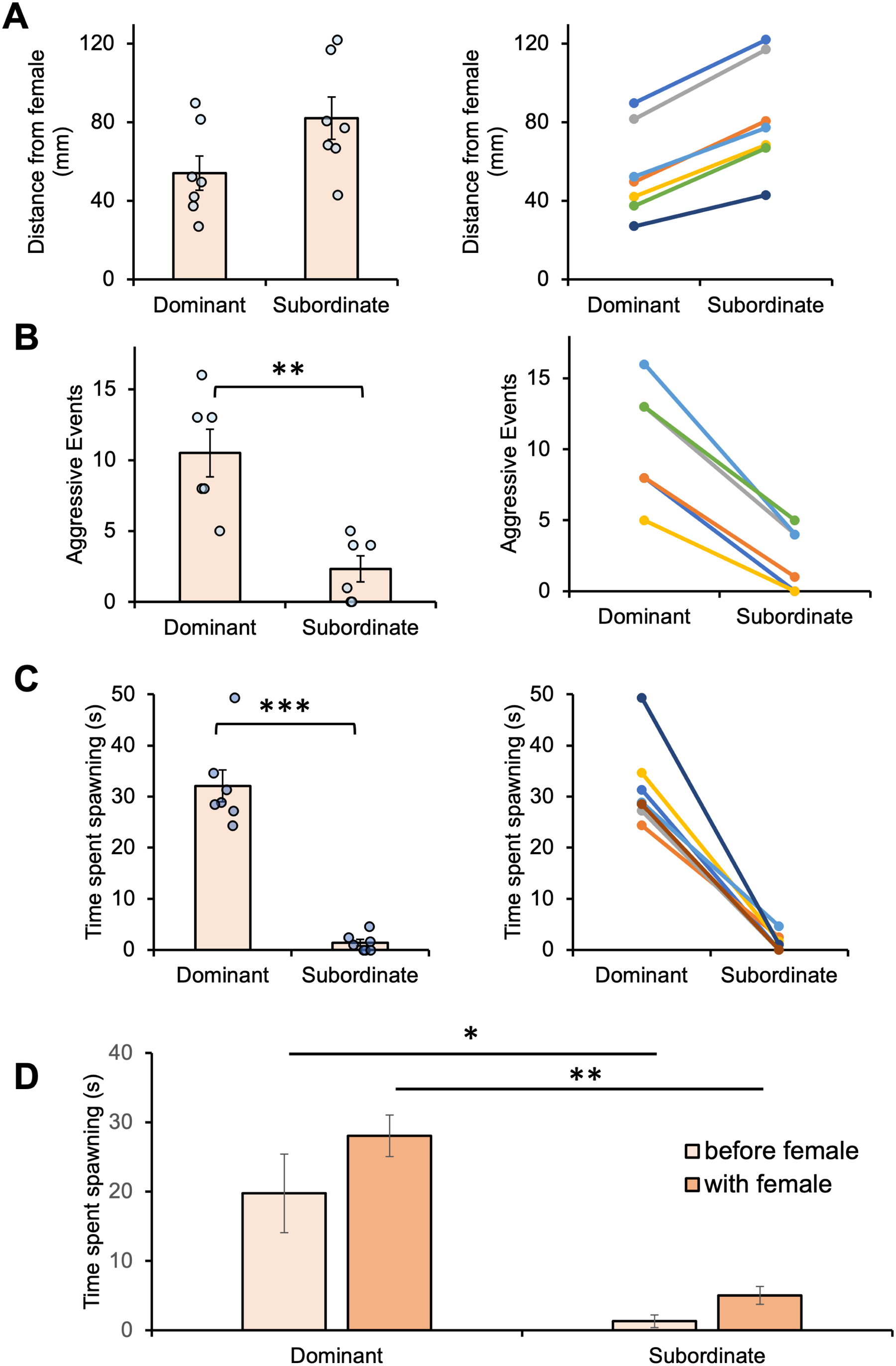
Dominance parameters in fish exposed to ALAN. **(A)** mate guarding behavior, measured as distance from the female (n=7), **(B)** aggression, counted as aggressive events (n=6), and **(C)** time spent spawning (n=7). The time spawning was then subdivided (D) in before and during female oviposition (n=10-11). Statistical analyses were performed using paired Student’s (A,C) and Welch’s test (B) t-tests, or Kruskal-Wallis test (D). The data are represented as mean ± SEM (left) and graphed according to paired competing males (right). Significant differences are indicated by asterisks (*:p< 0,05; **:p< 0,01; ***:p< 0,001).

In contrast, continuous light, which can be seen as an extreme ALAN condition, completely abolished the establishment of hierarchies. No aggressive events and no guarding behavior could be observed under these conditions as shown by the very similar distance between the two males and the female (figure 5A). In fact, other behaviors were also affected by continuous light. When measuring the time fish occupied the bottom, middle or top part of the tank (figure 5B), we found that fish exposed to continuous light spent significantly more time on the bottom and less on the top than fish exposed to control light conditions (figure 5C). Fish exposed to continuous light and ALAN also swam significantly shorter distances than the control fish (figure 5D). Finally, we found that while swimming activity in the morning did not vary with light treatment, fish exposed to continuous light spent significantly more time swimming, and at a higher velocity, during night than control fish (figure 5E,F). Together these results showing that fish exposed to continuous light are more active through the night while similarly active to the other groups during the day, suggest that physical exhaustion may explain why they spend more time on the bottom part of the tank and do not show guarding behavior or aggressivity. Interestingly for the time spent in each zone across the water column, time spent moving and velocity parameters, fish exposed to ALAN always showed values between those of control and continuous light exposed fish groups, supporting the view that fish behavior is also affected by ALAN, although to a lesser degree than by continuous light. In addition, the distance traveled was significantly reduced in fish exposed to ALAN compared to control fish. Together, these results suggest that ALAN also have effects on fish behavior. Because continuous light prevented determination of hierarchy, we did not investigate this environmental condition further.

**Figure 5:**
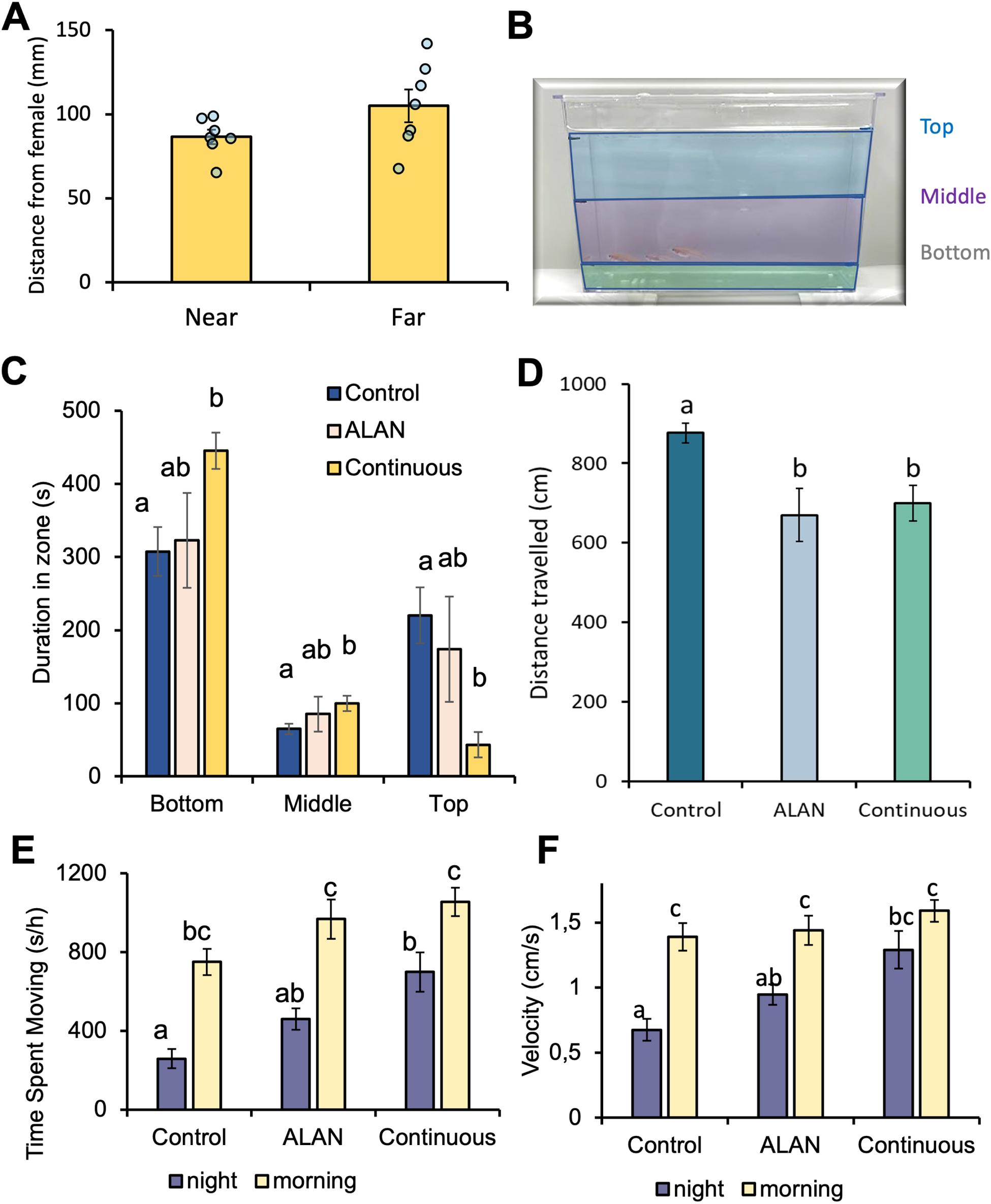
**(A)** Distance between the two males and the female when exposed to continuous light. We then did a water column test **(B)** to measure the time fish spend either in the bottom, middle or top part of the tank. **(C-D)** Graph showing the results of the water column test for fish kept sin control, ALAN or continuous light conditions (n=2). **(E-F)** Activity at night and during morning time of fish exposed to control light-dark cycle, ALAN, and continuous light (n=4). The data are represented as mean ± SEM. Statistical analyses were performed using two-way ANOVA with Tukey post-hoc and significant differences (p< 0.05) are indicated by different letters.

### Endocrine components do not differ between subordinate and dominant males neither in normal light conditions nor with light pollution at night

We looked for differences in the stress and reproductive endocrine systems between dominant and subordinate males. In the pituitary, mRNA levels for *lhb* and *fshb* did not differ between dominant and subordinate males, neither in control nor ALAN treated groups (figure 6A). Similarly, circulating levels of 11-ketotestoterone (11-KT) and testosterone (T) did not differ between dominant and control males with or without ALAN (figure 6B). Unexpectedly, we found no difference in circulating cortisol levels between dominant and subordinate males in any of the conditions (figure 6C). However, we observed that circulating cortisol levels varied widely among control males, but were more homogenous among males exposed to ALAN. Together these results suggest that neither pituitary nor gonadal hormones play a role in either the high fertilization rates of subordinate males despite their low access to females in control light regime, or the higher fertilization rates of dominant males in ALAN conditions.

**Figure 6:**
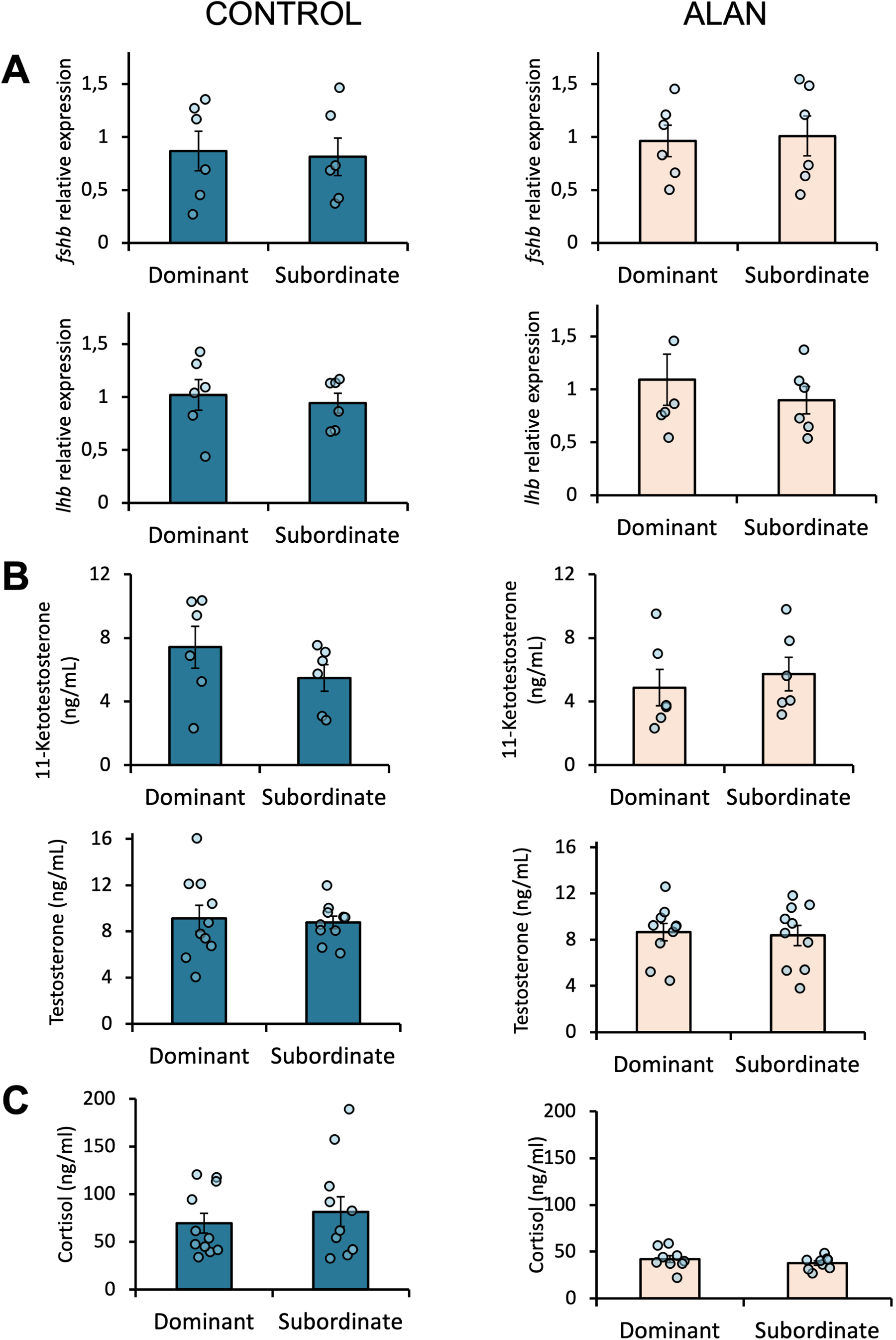
**(A)** Pituitary gonadotropin gene expression for dominant and subordinate males exposed to control (left) and artificial light at night (right). For both *fshb* and *lhb*, n=6. Circulating **(B)** sex steroid and **(C)** cortisol levels in dominant and subordinate males exposed to control (left) and artificial light at night (right). For 11-ketotestosterone, n=6, and for testosterone n=10. For cortisol, control n=11 and ALAN n=9. The data are represented as mean ± SEM. Differences between means were assessed using paired Student’s t-tests, and no differences were significant (p> 0.05).

### Light pollution affects sperm quality

Finally, we compared parameters of spermatogenesis in dominant and subordinate males in different light conditions. GSI, an index of gonadal development, did not significantly differ between dominant and subordinate males exposed to either control light or light pollution (figure 7A). Surprisingly, the percentage of motile sperm significantly increased in dominant males exposed to ALAN compared to control dominant males (figure 7B). The opposite seems to take place in subordinate males. Indeed, the percentage of motile sperm was reduced in subordinate males exposed to ALAN compared to control subordinate males, without being significantly different. However, our results show that while the percentage of motile sperm does not differ between dominant and subordinate males in control conditions, dominant males have a higher percentage of motile sperm than subordinate males when exposed to ALAN. Similar observations were made for the percentage of progressive sperm (figure 7C). Finally, we found that while the percentages of sperm moving at high, medium and low velocities were not different in subordinate and dominant males kept under control conditions, the percentage of both the high and medium velocity sperm is significantly higher in dominant males than in subordinate males exposed to ALAN (figure 7D). As high motility, progressivity, and velocity are characteristics of better sperm quality, these results may explain why dominant males exposed to ALAN fertilize more eggs than subordinate males.

**Figure 7:**
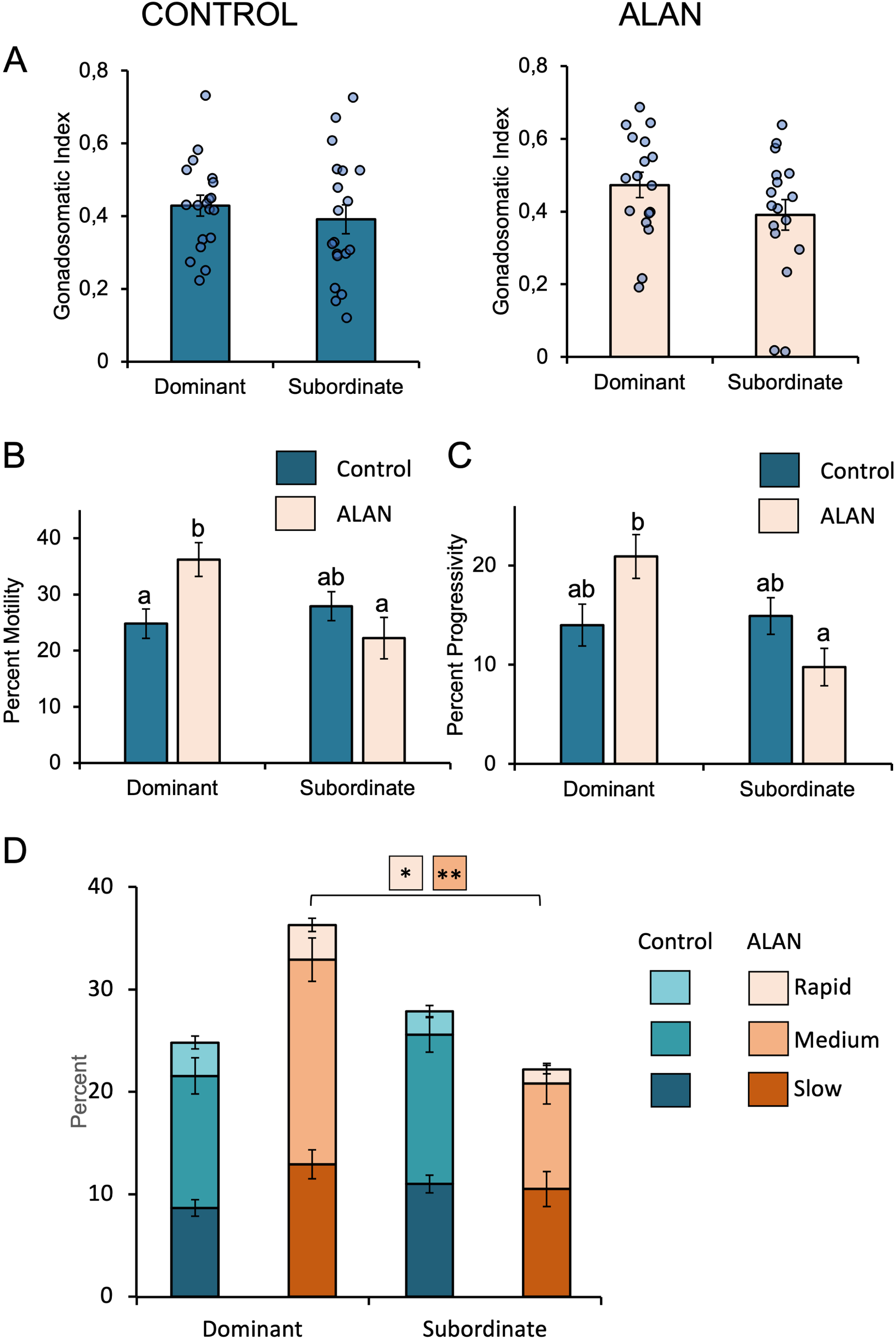
**(A)** Gonadosomatic index of dominant and subordinate males exposed to control light (n=19) and artificial light at night (n=17). Sperm quality of dominant and subordinate males indicated by **(B)** percentage of motile sperm, **(C)** percentage of progressive sperm, and **(D)** percentage of sperm moving at a rapid, medium, or slow velocity. For control fish, n=17; for fish exposed to ALAN, n=13. The data are represented as mean ± SEM. Statistical analyses were performed using ANOVAs with Tukey post-hoc and significant differences are indicated by different letters (p<0.05) or asterisks (* p<0.05; ** p<0.01).

## DISCUSSION

In the present study, we investigated the effect of the hierarchy status of male fish on their reproductive fitness, using the Japanese medaka as a model. A previous study showed that dominant male medaka display strong guarding behaviors when placed in a triadic relationship (two males with one females)(Yokoi et al., 2015). The same research group later suggested that dominant males, by preventing the subordinate male proximity to females, were increasing their fitness (Yokoi et al., 2016). However, another study reported sneaking behavior by subordinate males during spawning, allowing them to fertilize nearly half of the eggs (Weir, 2013). In the present study, we show that guarding behavior is not the only characteristic that can be used to decipher between dominant and subordinate males. Indeed, dominant males also displayed more aggression and spent more time spawning. Nevertheless, in agreement with the study from Weir and colleagues (Weir, 2013), we found that under normal light conditions subordinate males can produce half of the progeny despite their limited access to females. Our results also suggest that this success among subordinate males is linked to an efficient sneaking behavior; in contrast to dominant males which spend time spawning before the female releases eggs (premature spawning), the subordinate males spawned only during the critical time of egg release. It is interesting to see that while high male dominance rank has been associated with high reproductive success in several vertebrate species (e.g. red deer (*Cervus elaphus)* (Pemberton et al., 1992), northern elephant seals (*Mirounga angustirostris)* (Haley et al., 1994), African wild dogs (*Lycaon pictus)* (Girman et al., 1997), it is not always the case as shown in the chimpanzee (*Pan troglodytes schweinfurthii)* (Wroblewski et al., 2009) or in medaka fish with the present study.

Although we did not measure whether the dominant males released sperm during the premature spawning behavior, such release could explain why dominant males produce only 50% of the progeny. The high fertilization rate by subordinate males prompted us to investigate potential differences in sperm quality and in the endocrine factors regulating gametogenesis. However, we did not observe any difference in sperm quality between dominant and subordinate males, supporting the hypothesis that premature spawning leads to reduced sperm release during female oviposition. Curiously, we found no differences in expression levels for the two pituitary gonadotropins, GSI, or circulating T and 11-KT levels between subordinate and dominate males. This is quite different from what has been observed in *A. burtoni*, an African cichlid fish, in which higher plasma Lhb and Fshb protein levels, *lhb* and *fshb* mRNA levels, GSI (Maruska & Fernald, 2011), and sex steroid levels (Maruska & Fernald, 2013) were found in dominant compared to subordinate males. In addition, our results in medaka also differ from other fish studies where higher cortisol levels were observed in subordinate fish as for instance in the rainbow trout (*Oncorhynchus mykiss*) (Best & Gilmour, 2022; Doyon et al., 2003) and an African cichlid fish (*Haplochromis burtoni*) (Fox et al., 1997). While these results suggest species differences, these differences could also reflect the pronounced differences in the spawning cycle between fishes. While several teleost species spawn once per season, the medaka spawns daily during the breeding season. These differences could also result from different sampling times. Indeed, both sex steroids (Muhammad Rahmad Royan et al., 2023) and gonadotropin synthesis (Koger et al., 1999) show daily rhythms in medaka. Thus, the timing of the sampling might play a role in observing differences. It would be interesting to compare the daily rhythms of gonadotropins and sex steroids in dominant and subordinate males, in a future study.

We then investigated the effect of light pollution on medaka male fitness, as ALAN was previously reported to disrupt fish reproduction. ALAN was found to alter the behavior and activity smallmouth bass (*Micropterus dolomieu*) during its nest guarding period (Foster et al., 2016), potentially detracting from egg protection. ALAN has also been found to reduce gonadotropin and sex steroid levels, suggesting perturbation of seasonal reproduction in wild fish (Brüning, Hölker, et al., 2018; Brüning, Kloas, et al., 2018). In the present study, we found that the proportion of progeny produced by dominant and subordinate males changes drastically in presence of night light pollution, to the benefit of dominant males. This is to our knowledge the first evidence in any vertebrate that light pollution can differentially affect the fitness of specific individuals within a population. Whether the effects of light pollution observed in medaka can be extended to other teleost or vertebrate species remains to be determined, but if true, it would significantly impact the genetic pool of a population. Indeed, if dominants produce more progeny under conditions of nocturnal light pollution, their genotype will be selected. Decreased genetic diversity within a population might make the population more susceptible to disease and reduce the capacity of a population to adapt to a changing environment (Frankham et al., 2017; Hughes et al., 2008).

We then tried to identify which endocrine and behavioral parameters differed, between the dominant and subordinate males, that could explain the observed differences in reproductive success between male types when exposed to ALAN. While we did not observe any obvious change in the behavior, we found that sperm quality was affected by ALAN. Several indicators of sperm quality show that dominant males have higher quality sperm than subordinate males under such light conditions. Surprisingly, neither the GSI, gonadotropin expression, nor circulating sex steroid levels differed between subordinate and dominant males. This is surprising as the pituitary gonadotrope cell population is known to be very plastic, adapting the hormone production to the demand by increasing their activity and number (Romain Fontaine, Elia Ciani, et al., 2020). This plasticity has been clearly demonstrated in medaka (Fontaine et al., 2019; R. Fontaine et al., 2020). In addition, our recent work showed that reproduction and gonadotrope cell plasticity is highly dependent on photoperiod in medaka (M. R. Royan et al., 2023). Our finding that gonadotropins and sex steroids are not affected by light pollution in medaka also differs from previous studies in perch and roach (Brüning, Kloas, et al., 2018), in which ALAN decreased gonadotropin transcripts and circulating sex steroid levels. However, another study from the same research group found that gonadotropin transcript levels were not affected by ALAN in the roach (Brüning, Hölker, et al., 2018), suggesting that the effects on gonadotropins may be observed only under certain circumstances.

In conclusion, we found that light pollution can affect the fitness of medaka males within an established hierarchy by affecting the sperm quality. This has broad implications on fish ecology in both coastal marine and freshwater habitats where fish are exposed to anthropogenic light pollution (cities, harbors), with potential impacts on genetic diversity of these fish populations. The mechanistic link between light pollution and sperm quality remains to be determined.

## ACKNOWLEDGEMENTS

We thank Lourdes Carreon G Tan, Anthony Peltier and Dr. Arturas Kavaliauskis for their help with the maintenance of the fish.

## FUNDINGS

This work was supported by NMBU and Fulbright Norway.

## AUTHOR CONTRIBUTION

RF designed the experimental plan. LC did the experiments with the help from MRR, AS and RF, IM. LC draft the manuscript. RF and DB wrote the manuscript with the help from all authors.

---

**Supplemental figure 1:**
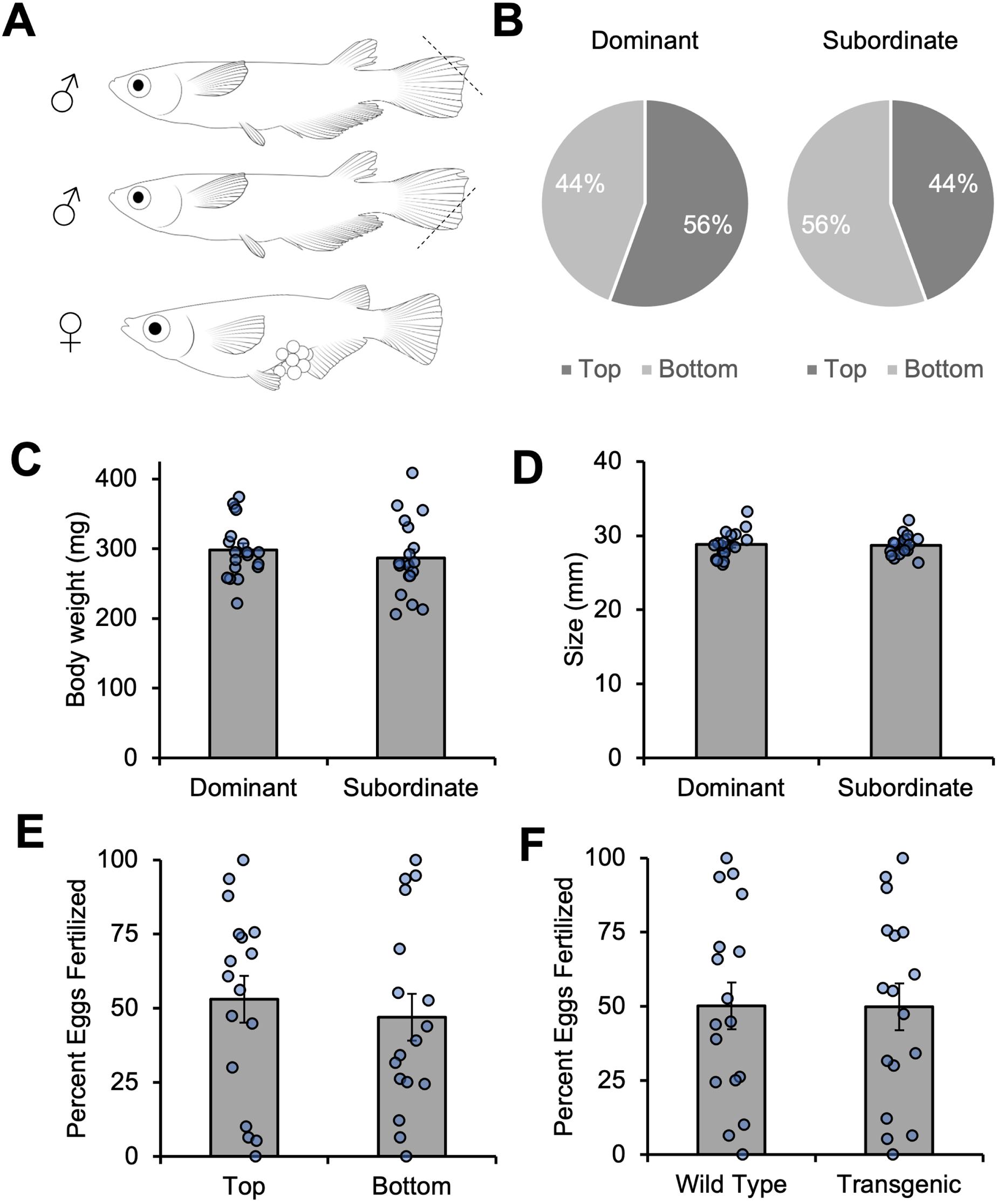
**(A)** Schematic drawing of the triadic setup used in this study in which each male was fin clipped on the top or bottom of the tail to allow tracking of individuals over time. **(B)** Graphics showing that the side where the tail finn was clipped did not affect the hierarchy status. **(C-D)** Graph showing that the body size and weight were kepts relatively similar between the males used in our experiments. **(E-F)** Graph showing that neuther the side where the tail finn was clipped not the genetic background affected the percentage of eggs males could fertilized.

**Supplemental figure 2:**
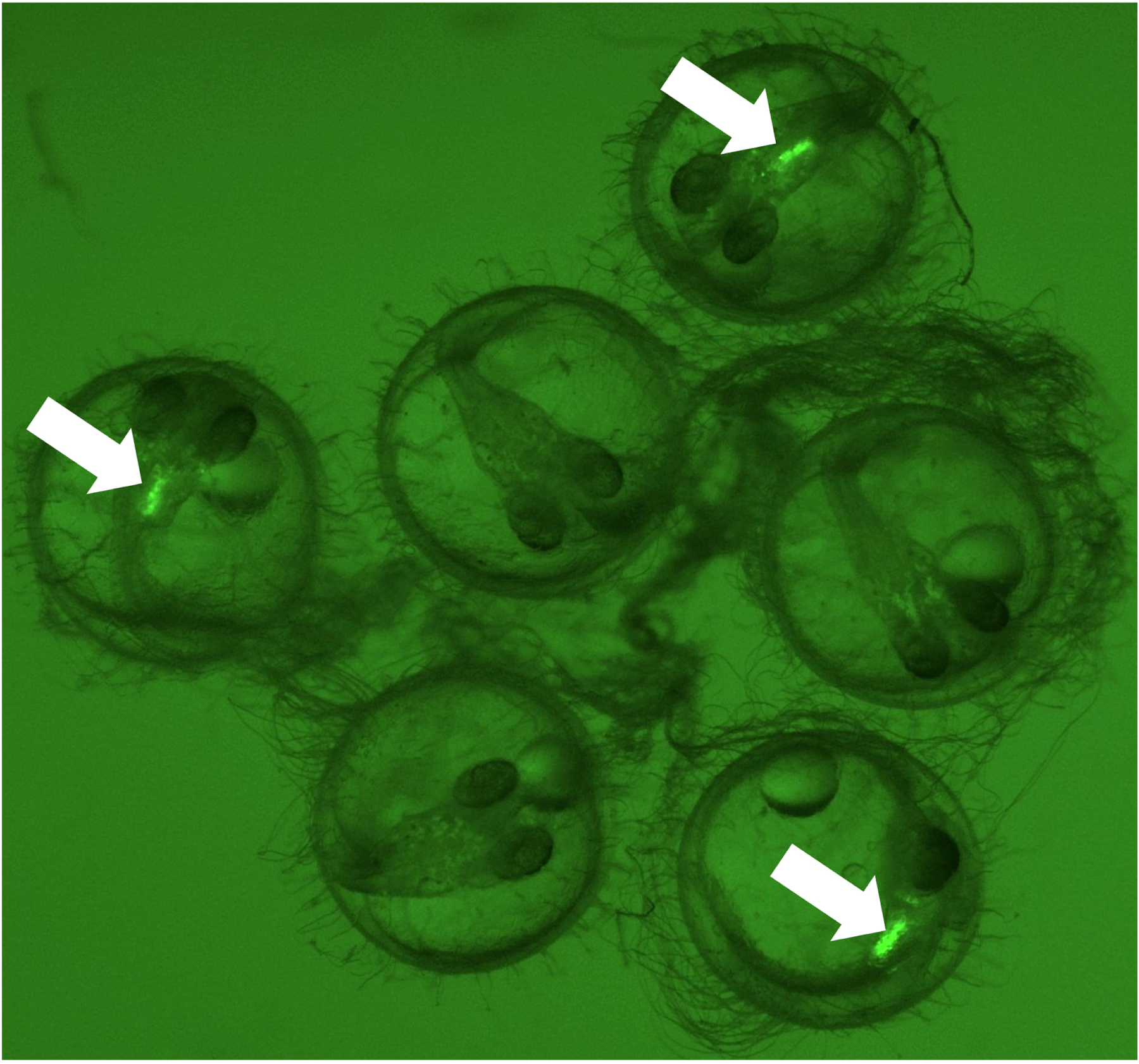
Picture of the transgenic lhb-hrGfpII embryos at 48 days post fertilization where the Gfp is easily identifiable (arrows) allowing us to assess male paternity.

## Notes

### Competing Interest Statement

The authors have declared no competing interest.

## REFERENCES

Ager-Wick, E., Hodne, K., Fontaine, R., Von Krogh, K., Haug, T. M., & Weltzien, F. A. (2018). Preparation of a High-quality Primary Cell Culture from Fish Pituitaries. Journal of Visualized Experiments(138). https://doi.org/10.3791/58159

Alberts, S. C., Buchan, J. C., & Altmann, J. (2006). Sexual selection in wild baboons: from mating opportunities to paternity success. Animal Behaviour, 72(5), 1177–1196. https://doi.org/https://doi.org/10.1016/j.anbehav.2006.05.001

Altmann, J., Sapolsky, R., & Licht, P. (1995). Baboon fertility and social status. Nature, 377(6551), 688–690. https://doi.org/10.1038/377688a0

Bennett, N. C. (1994). Reproductive suppression in social Cryptomys damarensis colonies—a lifetime of socially-induced sterility in males and females (Rodentia: Bathyergidae) [https://doi.org/10.1111/j.1469-7998.1994.tb06054.x]. Journal of Zoology, 234(1), 25–39. https://doi.org/https://doi.org/10.1111/j.1469-7998.1994.tb06054.x

Best, C., & Gilmour, K. M. (2022). Regulation of cortisol production during chronic social stress in rainbow trout. General and Comparative Endocrinology, 325, 114056. https://doi.org/https://doi.org/10.1016/j.ygcen.2022.114056

Brüning, A., Hölker, F., Franke, S., Kleiner, W., & Kloas, W. (2018). Influence of light intensity and spectral composition of artificial light at night on melatonin rhythm and mRNA expression of gonadotropins in roach Rutilus rutilus. Fish Physiol Biochem, 44(1), 1–12. https://doi.org/10.1007/s10695-017-0408-6

Brüning, A., Kloas, W., Preuer, T., & Hölker, F. (2018). Influence of artificially induced light pollution on the hormone system of two common fish species, perch and roach, in a rural habitat. Conserv Physiol, 6(1), coy016. https://doi.org/10.1093/conphys/coy016

Burow, S., Fontaine, R., von Krogh, K., Mayer, I., Nourizadeh-Lillabadi, R., Hollander-Cohen, L., Cohen, Y., Shpilman, M., Levavi-Sivan, B., & Weltzien, F.-A. (2019). Medaka follicle-stimulating hormone (Fsh) and luteinizing hormone (Lh): Developmental profiles of pituitary protein and gene expression levels. General and Comparative Endocrinology, 272, 93–108. https://doi.org/https://doi.org/10.1016/j.ygcen.2018.12.006

Closs, E. L., Royan Muhammad, R., Sayyari, A., Mayer, I., Weltzien, F.-A., Baker, D. M., & Fontaine, R. (2023). Impact of light pollution at night on male reproductive success in Japanese medaka (Oryzias latipes) Version DRAFT VERSION) DataverseNO. https://doi.org/doi:10.18710/NZJVB7

Closs, L., Sayarri, A., & Fontaine, R. (2022). Sperm Collection and Computer-Assisted Sperm Analysis in the Teleost Model Japanese Medaka (Oryzias latipes). Journal of Visualized Experiments. https://doi.org/10.3791/64326

Davies, T. W., Duffy, J. P., Bennie, J., & Gaston, K. J. (2014). The nature, extent, and ecological implications of marine light pollution [https://doi.org/10.1890/130281]. Frontiers in Ecology and the Environment, 12(6), 347–355. https://doi.org/https://doi.org/10.1890/130281

Doyon, C., Gilmour, K. M., Trudeau, V. L., & Moon, T. W. (2003). Corticotropin-releasing factor and neuropeptide Y mRNA levels are elevated in the preoptic area of socially subordinate rainbow trout. General and Comparative Endocrinology, 133(2), 260–271. https://doi.org/https://doi.org/10.1016/S0016-6480(03)00195-3

Dufour, S., Sebert, M. E., Weltzien, F. A., Rousseau, K., & Pasqualini, C. (2010). Neuroendocrine control by dopamine of teleost reproduction. J Fish Biol, 76(1), 129–160. https://doi.org/10.1111/j.1095-8649.2009.02499.x

Endal, H. P., Taranger, G. L., Stefansson, S. O., & Hansen, T. (2000). Effects of continuous additional light on growth and sexual maturity in Atlantic salmon, Salmo salar, reared in sea cages. Aquaculture, 191(4), 337–349. https://doi.org/10.1016/S0044-8486(00)00444-0

Fernald, R. D., & Hirata, N. R. (1977). Field study of Haplochromis burtoni: Quantitative behavioural observations. Animal Behaviour, 25, 964–975. https://doi.org/https://doi.org/10.1016/0003-3472(77)90048-3

Fjelldal, P. G., Schulz, R. W., Nilsen, T. O., Andersson, E., Norberg, B., & Hansen, T. J. (2018). Sexual maturation and smoltification in domesticated Atlantic salmon (Salmo salar L.) – is there a developmental conflict? Physiological Reports, 6(17), e13809–e13809. https://doi.org/10.14814/phy2.13809

Fontaine, R., Ager-Wick, E., Hodne, K., & Weltzien, F.-A. (2019). Plasticity of Lh cells caused by cell proliferation and recruitment of existing cells. Journal of Endocrinology, 240(2), 361–377. https://doi.org/http://doi.org/10.1530/JOE-18-0412

Fontaine, R., Ager-Wick, E., Hodne, K., & Weltzien, F. A. (2020). Plasticity in medaka gonadotropes via cell proliferation and phenotypic conversion. J Endocrinol, 245(1), 21–37. https://doi.org/10.1530/JOE-19-0405

Fontaine, R., Ciani, E., Haug, T. M., Hodne, K., Ager-Wick, E., Baker, D. M., & Weltzien, F.-A. (2020). Gonadotrope plasticity at cellular, population and structural levels: A comparison between fishes and mammals. General and Comparative Endocrinology, 287, 113344. https://doi.org/10.1016/j.ygcen.2019.113344

Fontaine, R., Royan, M. R., von Krogh, K., Weltzien, F.-A., & Baker, D. M. (2020). Direct and Indirect Effects of Sex Steroids on Gonadotrope Cell Plasticity in the Teleost Fish Pituitary [Review]. Frontiers in Endocrinology, 11(858). https://doi.org/10.3389/fendo.2020.605068

Foster, J. G., Algera, D. A., Brownscombe, J. W., Zolderdo, A. J., & Cooke, S. J. (2016). Consequences of Different Types of Littoral Zone Light Pollution on the Parental Care Behaviour of a Freshwater Teleost Fish. Water, Air, & Soil Pollution, 227(11), 404. https://doi.org/10.1007/s11270-016-3106-6

Fox, H. E., White, S. A., Kao, M. H., & Fernald, R. D. (1997). Stress and dominance in a social fish. J Neurosci, 17(16), 6463–6469. https://doi.org/10.1523/jneurosci.17-16-06463.1997

Frankham, R., Ballou, J. D., Ralls, K., Eldridge, M. D. B., Dudash, M. R., Fenster, C. B., Lacy, R. C., & Sunnucks, P. (2017). 65Loss of genetic diversity reduces ability to adapt. In R. Frankham, J. D. Ballou, K. Ralls, M. Eldridge, M. R. Dudash, C. B. Fenster, R. C. Lacy, & P. Sunnucks (Eds.), Genetic Management of Fragmented Animal and Plant Populations (pp. 0). Oxford University Press. https://doi.org/10.1093/oso/9780198783398.003.0004

Fukamachi, S., Kinoshita, M., Aizawa, K., Oda, S., Meyer, A., & Mitani, H. (2009). Dual control by a single gene of secondary sexual characters and mating preferences in medaka. BMC Biol, 7, 64. https://doi.org/10.1186/1741-7007-7-641741-7007-7-64 [pii]

Girman, D. J., Mills, M., Geffen, E., & Wayne, R. K. (1997). A molecular genetic analysis of social structure, dispersal, and interpack relationships of the African wild dog (Lycaon pictus). Behavioral Ecology and Sociobiology, 40, 187–198.

Haley, M. P., Deutsch, C. J., & Le Boeuf, B. J. (1994). Size, dominance and copulatory success in male northern elephant seals, Mirounga angustirostris. Animal Behaviour, 48(6), 1249–1260.

Hildahl, J., Sandvik, G. K., Lifjeld, R., Hodne, K., Nagahama, Y., Haug, T. M., Okubo, K., & Weltzien, F. A. (2012). Developmental tracing of luteinizing hormone beta-subunit gene expression using green fluorescent protein transgenic medaka (Oryzias latipes) reveals a putative novel developmental function. Developmental Dynamics, 241(11), 1665–1677. https://doi.org/10.1002/dvdy.23860

Hodne, K., Fontaine, R., Ager-Wick, E., & Weltzien, F. A. (2019). Gnrh1-Induced Responses Are Indirect in Female Medaka Fsh Cells, Generated Through Cellular Networks. Endocrinology, 160(12), 3018–3032. https://doi.org/10.1210/en.2019-00595

Hughes, A. R., Inouye, B. D., Johnson, M. T. J., Underwood, N., & Vellend, M. (2008). Ecological consequences of genetic diversity [https://doi.org/10.1111/j.1461-0248.2008.01179.x]. Ecology Letters, 11(6), 609–623. https://doi.org/https://doi.org/10.1111/j.1461-0248.2008.01179.x

Kasahara, M., Naruse, K., Sasaki, S., Nakatani, Y., Qu, W., Ahsan, B., Yamada, T., Nagayasu, Y., Doi, K., Kasai, Y., Jindo, T., Kobayashi, D., Shimada, A., Toyoda, A., Kuroki, Y., Fujiyama, A., Sasaki, T., Shimizu, A., Asakawa, S., … Kohara, Y. (2007). The medaka draft genome and insights into vertebrate genome evolution. Nature, 447(7145), 714–719. https://doi.org/http://doi.org/10.1038/nature05846

Kinoshita, M., Murata, K., Naruse, K., & Tanaka, M. (2009). Looking at Adult Medaka. In K. M. M. Kinoshita, K. Naruse & M. Tanaka (Ed.), Medaka: Biology, Management, and Experimental Protocols (pp. 117–164). Wiley Online Library. https://books.google.it/books?hl=it&lr=&id=z6JBkqBNURQC&oi=fnd&pg=PR15&dq=Kinoshita+M,+Murata+K,+Naruse+K,+Tanaka+M.,+2009.+Medaka:+biology,+management,+and+experimental+protocols.+1st+ed.+Ames,+IA:+Wiley-Blackwell.&ots=4hVWv5X1uZ&sig=mGuRGbSHeyIGniRjGxdI

Kobayashi, M., Yoritsune, T., Suzuki, S., Shimizu, A., Koido, M., Kawaguchi, Y., Hayakawa, Y., Eguchi, S., Yokota, H., & Yamamoto, Y. (2012). Reproductive behavior of wild medaka in an outdoor pond. Nippon Suisan Gakkaishi, 78(5), 922–933.

Koger, C. S., Teh, S. J., & Hinton, D. E. (1999). Variations of Light and Temperature Regimes and Resulting Effects on Reproductive Parameters in Medaka (Oryzias latipes)1. Biology of Reproduction, 61(5), 1287–1293. https://doi.org/10.1095/biolreprod61.5.1287

Koyama, S., & Kamimura, S. (2000). Influence of social dominance and female odor on the sperm activity of male mice. Physiol Behav, 71(3-4), 415–422. https://doi.org/10.1016/s0031-9384(00)00361-9

Kummu, M., Moel, H., Ward, P., & Olli, V. (2011). How Close Do We Live to Water? A Global Analysis of Population Distance to Freshwater Bodies. PLoS One, 6, e20578. https://doi.org/10.1371/journal.pone.0020578

Kyba, C. C. M., Kuester, T., Sánchez de Miguel, A., Baugh, K., Jechow, A., Hölker, F., Bennie, J., Elvidge, C. D., Gaston, K. J., & Guanter, L. (2017). Artificially lit surface of Earth at night increasing in radiance and extent. Science Advances, 3(11), e1701528. https://doi.org/10.1126/sciadv.1701528

Lucassen, E. A., Coomans, C. P., van Putten, M., de Kreij, S. R., van Genugten, J. H., Sutorius, R. P., de Rooij, K. E., van der Velde, M., Verhoeve, S. L., Smit, J. W., Löwik, C. W., Smits, H. H., Guigas, B., Aartsma-Rus, A. M., & Meijer, J. H. (2016). Environmental 24-hr Cycles Are Essential for Health. Curr Biol, 26(14), 1843–1853. https://doi.org/10.1016/j.cub.2016.05.038

Marangoni, L. F. B., Davies, T., Smyth, T., Rodríguez, A., Hamann, M., Duarte, C., Pendoley, K., Berge, J., Maggi, E., & Levy, O. (2022). Impacts of artificial light at night in marine ecosystems—A review [https://doi.org/10.1111/gcb.16264]. Global Change Biology, 28(18), 5346–5367. https://doi.org/https://doi.org/10.1111/gcb.16264

Maruska, K. P., & Fernald, R. D. (2011). Plasticity of the reproductive axis caused by social status change in an african cichlid fish: II. testicular gene expression and spermatogenesis. Endocrinology, 152(1), 291–302. https://doi.org/10.1210/en.2010-0876

Maruska, K. P., & Fernald, R. D. (2013). Social regulation of male reproductive plasticity in an African cichlid fish. Integr Comp Biol, 53(6), 938–950. https://doi.org/10.1093/icb/ict017

Matsuda, M. (2005). Sex determination in the teleost medaka, Oryzias latipes. Annu Rev Genet, 39, 293–307. https://doi.org/10.1146/annurev.genet.39.110304.095800

Munakata, A., & Kobayashi, M. (2009). Endocrine control of sexual behavior in teleost fish. General and Comparative Endocrinology, 165, 456–468. https://doi.org/10.1016/j.ygcen.2009.04.011

Nanda, I., Kondo, M., Hornung, U., Asakawa, S., Winkler, C., Shimizu, A., Shan, Z., Haaf, T., Shimizu, N., Shima, A., Schmid, M., & Schartl, M. (2002). A duplicated copy of DMRT1 in the sex-determining region of the Y chromosome of the medaka, Oryzias latipes. Proc Natl Acad Sci U S A, 99(18), 11778–11783. https://doi.org/10.1073/pnas.182314699

Navara, K. J., & Nelson, R. J. (2007). The dark side of light at night: physiological, epidemiological, and ecological consequences. J Pineal Res, 43(3), 215–224. https://doi.org/10.1111/j.1600-079X.2007.00473.x

O’Connor, J. J., Fobert, E. K., Besson, M., Jacob, H., & Lecchini, D. (2019). Live fast, die young: Behavioural and physiological impacts of light pollution on a marine fish during larval recruitment. Mar Pollut Bull, 146, 908–914. https://doi.org/10.1016/j.marpolbul.2019.05.038

Ono, Y., & Uematsu, T. (1957). Mating Ethogram in Oryzias latipes (With 1 Text-figure). 北海道大學理學部紀要= JOURNAL OF THE FACULTY OF SCIENCE HOKKAIDO UNIVERSITY Series ⅤⅠ. ZOOLOGY, 13(1-4), 197–202.

Oppedal, F., Berg, A., Olsen, R. E., Taranger, G. L., & Hansen, T. (2006). Photoperiod in seawater influence seasonal growth and chemical composition in autumn sea-transferred Atlantic salmon (Salmo salar L.) given two vaccines. Aquaculture, 254(1), 396–410. https://doi.org/https://doi.org/10.1016/j.aquaculture.2005.10.026

Pemberton, J., Albon, S., Guinness, F., Clutton-Brock, T., & Dover, G. (1992). Behavioral estimates of male mating success tested by DNA fingerprinting in a polygynous mammal. Behavioral Ecology, 3(1), 66–75.

Pulgar, J., Zeballos, D., Vargas, J., Aldana, M., Manriquez, P. H., Manriquez, K., Quijón, P. A., Widdicombe, S., Anguita, C., Quintanilla, D., & Duarte, C. (2019). Endogenous cycles, activity patterns and energy expenditure of an intertidal fish is modified by artificial light pollution at night (ALAN). Environmental Pollution, 244, 361–366. https://doi.org/https://doi.org/10.1016/j.envpol.2018.10.063

Rhees, R. W., Kirk, B. A., Sephton, S., & Lephart, E. D. (1997). Effects of Prenatal Testosterone on Sexual Behavior, Reproductive Morphology and LH Secretion in the Female Rat. Developmental Neuroscience, 19(5), 430–437. https://doi.org/10.1159/000111240

Rowan, W. (1938). LIGHT AND SEASONAL REPRODUCTION IN ANIMALS [https://doi.org/10.1111/j.1469-185X.1938.tb00523.x]. Biological Reviews, 13(4), 374–401. https://doi.org/https://doi.org/10.1111/j.1469-185X.1938.tb00523.x

Royan, M. R., Hodne, K., Rasoul, N. L., Henkel, C. V., Weltzien, F.-A., & Fontaine, R. (2023). Photoperiod regulates gonadotrope cell mitosis in medaka via melatonin, Tsh and folliculostellate cells. in prep.

Royan, M. R., Kanda, S., Kayo, D., Song, W., Ge, W., Weltzien, F.-A., & Fontaine, R. (2020). Gonadectomy and blood sampling procedures in the small size teleost model Japanese medaka (Oryzias latipes). JoVE. https://doi.org/https://doi.org/10.1101/2020.08.30.271478

Royan, M. R., Kayo, D., Weltzien, F.-A., & Fontaine, R. (2023). Sexually Dimorphic Regulation of Gonadotrope Cell Hyperplasia in Medaka Pituitary via Mitosis and Transdifferentiation. Endocrinology, 164(4), bqad030. https://doi.org/10.1210/endocr/bqad030

Royan, M. R., Siddique, K., Csucs, G., Puchades, M. A., Nourizadeh-Lillabadi, R., Bjaalie, J. G., Henkel, C. V., Weltzien, F.-A., & Fontaine, R. (2021). 3D Atlas of the Pituitary Gland of the Model Fish Medaka (Oryzias latipes) [Original Research]. Frontiers in Endocrinology, 12(912). https://doi.org/https://doi.org/10.3389/fendo.2021.719843

Schroer, S., & Hölker, F. (2017). Impact of Lighting on Flora and Fauna. In R. Karlicek, C.-C. Sun, G. Zissis, & R. Ma (Eds.), Handbook of Advanced Lighting Technology (pp. 957–989). Springer International Publishing. https://doi.org/10.1007/978-3-319-00176-0_42

Shima, A., & Mitani, H. (2004). Medaka as a research organism: past, present and future. Mechanisms of Development, 121(7–8), 599–604. https://doi.org/http://dx.doi.org/10.1016/j.mod.2004.03.011

Stefansson, S. O., Bjömsson, B. T., Hansen, T., Haux, C., Taranger, G. L., & Saunders, R. L. (1991). Growth, Parr–Smolt Transformation, and Changes in Growth Hormone of Atlantic Salmon (Salmo salar) Reared under Different Photoperiods. Canadian Journal of Fisheries and Aquatic Sciences, 48(11), 2100–2108. https://doi.org/10.1139/f91-249

Walker II, W. H., Meléndez-Fernández, O. H., Nelson, R. J., & Reiter, R. J. (2019). Global climate change and invariable photoperiods: A mismatch that jeopardizes animal fitness. Ecology and Evolution, 9(17), 10044–10054. https://doi.org/https://doi.org/10.1002/ece3.5537

Weir, L. (2013). Male–male competition and alternative male mating tactics influence female behavior and fertility in Japanese medaka (Oryzias latipes). Behavioral Ecology and Sociobiology, 67, 193. https://doi.org/10.1007/s00265-012-1438-9

Weltzien, F. A., Andersson, E., Andersen, O., Shalchian-Tabrizi, K., & Norberg, B. (2004). The brain-pituitary-gonad axis in male teleosts, with special emphasis on flatfish (Pleuronectiformes). Comp Biochem Physiol A Mol Integr Physiol, 137(3), 447–477. https://doi.org/10.1016/j.cbpb.2003.11.007

Wittbrodt, J., Shima, A., & Schartl, M. (2002). Medaka--a model organism from the far East. Nat Rev Genet, 3(1), 53–64. https://doi.org/10.1038/nrg704

Wroblewski, E. E., Murray, C. M., Keele, B. F., Schumacher-Stankey, J. C., Hahn, B. H., & Pusey, A. E. (2009). Male dominance rank and reproductive success in chimpanzees, Pan troglodytes schweinfurthii. Anim Behav, 77(4), 873–885. https://doi.org/10.1016/j.anbehav.2008.12.014

Yaron, Z., Gur, G., Melamed, P., Rosenfeld, H., Elizur, A., & Levavi-Sivan, B. (2003). Regulation of fish gonadotropins. Int Rev Cytol, 225, 131–185. http://www.ncbi.nlm.nih.gov/pubmed/12696592

Yokoi, S., Ansai, S., Kinoshita, M., Naruse, K., Kamei, Y., Young, L. J., Okuyama, T., & Takeuchi, H. (2016). Mate-guarding behavior enhances male reproductive success via familiarization with mating partners in medaka fish. Front Zool, 13, 21. https://doi.org/10.1186/s12983-016-0152-2

Yokoi, S., Okuyama, T., Kamei, Y., Naruse, K., Taniguchi, Y., Ansai, S., Kinoshita, M., Young, L. J., Takemori, N., Kubo, T., & Takeuchi, H. (2015). An essential role of the arginine vasotocin system in mate-guarding behaviors in triadic relationships of medaka fish (Oryzias latipes). PLoS Genet, 11(2), e1005009. https://doi.org/10.1371/journal.pgen.1005009

Zohar, Y., Munoz-Cueto, J. A., Elizur, A., & Kah, O. (2010). Neuroendocrinology of reproduction in teleost fish. Gen Comp Endocrinol, 165(3), 438–455. https://doi.org/10.1016/j.ygcen.2009.04.017

